# Frequency and complexity of de novo structural mutation in autism

**DOI:** 10.1101/030270

**Authors:** William M Brandler, Danny Antaki, Madhusudan Gujral, Amina Noor, Gabriel Rosanio, Timothy R Chapman, Daniel J Barrera, Guan Ning Lin, Dheeraj Malhotra, Amanda C Watts, Lawrence C Wong, Jasper A Estabillo, Therese E Gadomski, Oanh Hong, Karin V Fuentes Fajardo, Abhishek Bhandari, Renius Owen, Michael Baughn, Jeffrey Yuan, Terry Solomon, Alexandra G Moyzis, Stephan J Sanders, Gail E Reiner, Keith K Vaux, Charles M Strom, Kang Zhang, Alysson R Muotri, Natacha Akshoomoff, Suzanne M Leal, Karen Pierce, Eric Courchesne, Lilia M Iakoucheva, Christina Corsello, Jonathan Sebat

**Author notes:** Equal contributor (William M Brandler, Danny Antaki, Madhusudan Gujral).

## Abstract

Genetic studies of Autism Spectrum Disorder (ASD) have established that *de novo* duplications and deletions contribute to risk. However, ascertainment of structural variation (SV) has been restricted by the coarse resolution of current approaches. By applying a custom pipeline for SV discovery, genotyping and *de novo* assembly to genome sequencing of 235 subjects, 71 cases, 26 sibling controls and their parents, we present an atlas of 1.2 million SVs (5,213/genome), comprising 11 different classes. We demonstrate a high diversity of *de novo* mutations, a majority of which were undetectable by previous methods. In addition, we observe complex mutation clusters where combinations of *de novo* SVs, nucleotide substitutions and indels occurred as a single event. We estimate a high rate of structural mutation in humans (20%). Genetic risk for ASD is attributable to an elevated frequency of gene-disrupting *de novo* SVs but not an elevated rate of genome rearrangement.

## Introduction

Structural variants (SVs) such as deletions and duplications are a major source of genetic differences between humans and a significant contributor to risk for common disease [1]. In particular, studies of copy number variation (CNV) have been seminal in establishing a role for rare genetic variants in the etiology of Autism Spectrum Disorders (ASDs) [2, 3]. Despite this success, characterization of SVs from patient genomes remains a major challenge. Identification of SVs in human populations and disease have been restricted by the limited sensitivity of microarray- and sequencing-based approaches [4, 5, 6].

Large CNVs detectable by microarrays represent a small fraction of structural variation in the genome. Recent methodological advances have enabled the discovery of a wide variety of SV classes from whole genome sequencing (WGS) datasets, including small deletions and duplications down to 50 base pairs (bp) in length, inversions, translocations, mobile element insertions, and more complex rearrangements. By applying a combination of specialized methods, each tailored to specific classes of variation, the 1000 Genomes Project (1KG) has produced the most complete catalogue of SVs to date, creating an integrated call set of eight classes of SV using low-coverage (7.4×) WGS in 2,504 human genomes [7]. In a study of 250 population-control families, analysis of low coverage (13×) WGS data allowed for detection of *de novo* deletions, tandem duplications, and mobile element insertions [8]. However, advanced analytical methods for SV discovery and genotyping have not been applied in genetic studies of ASD or intellectual disability. Initial forays into the application of WGS to detection of SVs in neurodevelopmental disorders have been either restricted to CNVs larger than 1kb [9], focused on a subset of variant calls prioritized by putative clinical-relevance [10, 9], or limited to the characterization of CNVs previously detected using microarrays [4].

More comprehensive ascertainment of SV is needed to elucidate the genetic mechanisms that underlie risk for ASD. Here we apply a suite of complementary SV discovery methods coupled with novel methods for SV genotyping and *de novo* mutation detection, to assess global patterns and rates of structural mutation in ASD.

## Results

### Genome Sequencing Uncovers a Diverse Landscape of Structural Variation

We recruited patients with ASD and their families from Rady Children’s Hospital San Diego and local pediatric clinics, including 235 subjects, 71 cases and 26 typically developing sibling controls. WGS of blood-derived genomic DNA was performed at a mean coverage of 40.6× (Table S1; see Methods).

We developed a SV discovery pipeline that utilizes a combination of three specialized methods that are each optimized to capture a specific subtype of variation. (**Figure S1** and Methods): (1) ForestSV [11] is a statistical learning approach we developed that integrates a variety of features from WGS data into a random forest classifier, and is optimized for the detection of deletions and duplications; (2) Lumpy [12] utilizes information from discordant paired-ends and split reads, and is optimal for the detection of balanced rearrangements such as inversions and translocations; (3) Mobster [13] utilizes paired-end and split reads for the detection of mobile element insertions (MEIs). As we show here, this combination of methods is highly efficient and provides accurate detection of most known classes of SV. For each subject, unfiltered SV calls from three methods were merged into a set of consensus calls (see methods).

The final call set from 235 subjects includes 1,225,067 SVs (5,213/genome) from 29,719 sites (**Figure 1**). The primary variant calls comprised seven major classes, including deletions (3,383 alleles / individual; 18,359 sites), duplications (423 / individual; 2,360 sites), inversions (51 / individual; 211 sites), and four classes of MEI (1,105 / individual; 7,915 sites) (**Figure 1; Supplementary Data Set 1** and **Table S2**). False discovery rates (FDR) for deletion and duplication calls were estimated from Illumina 2.5M SNP array data (using SVtoolkit, see methods), which was collected on a majority of samples in our study (*n* = 205). FDR was determined to be 7.0% for deletions and 9.2% for duplications. The complete call set and detailed descriptive information for all calls is provided in **Supplementary Data Set 1**. A comparison of our SV call set with the phase 3 SV call set from the 1KG is described in **Figure S2**.

**Figure 1.**
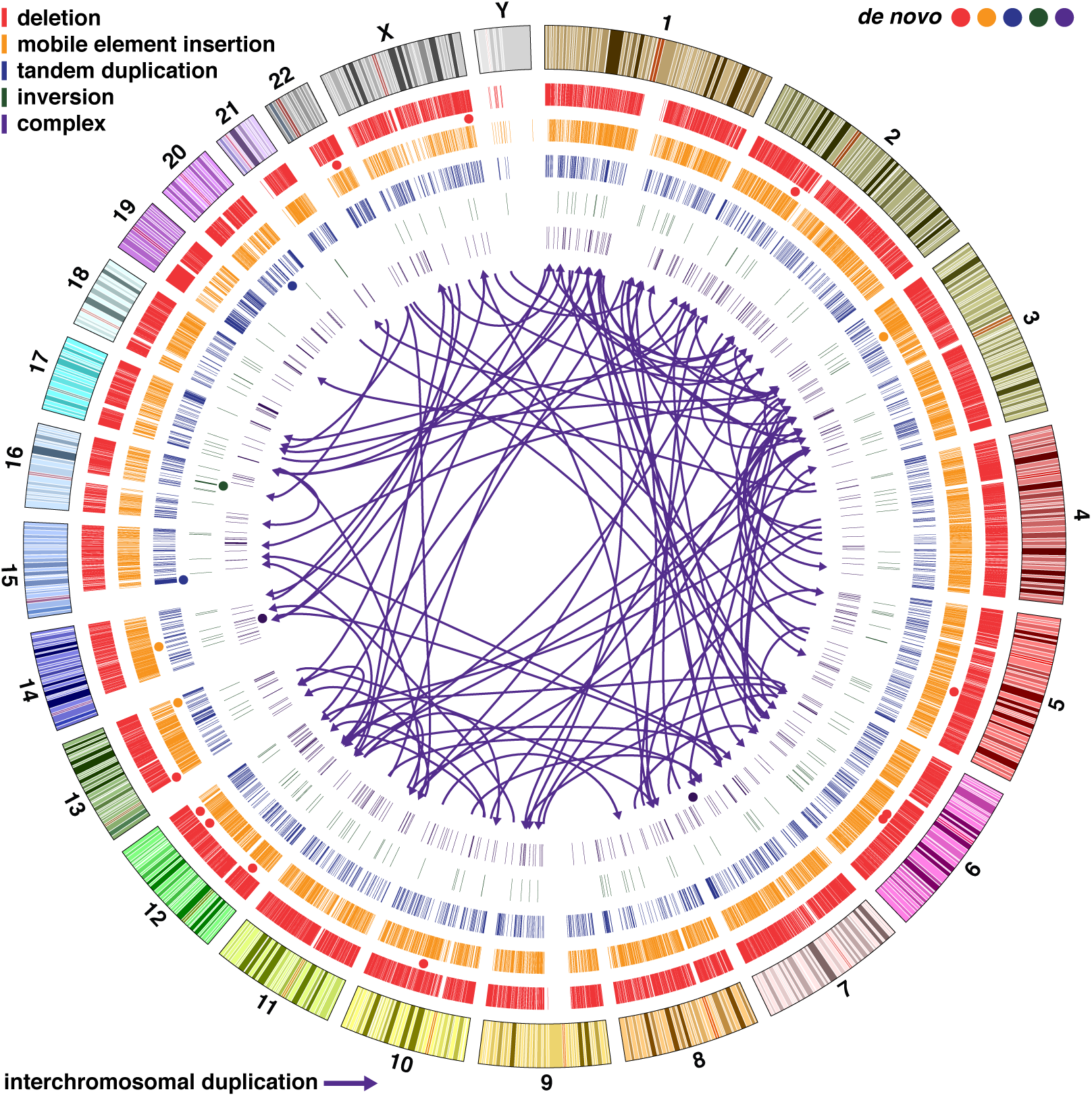
Structural Variation Detected from Whole Genome Sequencing in 235 Individuals. Circos plot with concentric circles representing (from outermost to inner): ideogram of the human genome with colored karyotype bands (hg19), deletions, mobile element insertions (four different classes), tandem duplications, balanced inversions, complex structural variants (four different classes). Circles indicate the location of *de novo* SVs, and their colors match the five SV types. Arrows represent interchromosomal duplications.

### Detection of Complex Rearrangements

A recent study using a combination of microarrays and sequencing of large-insert (‘jumping’) libraries has shown that a variety of complex structural variants are observed in a subset (24%) of ASD [4]. In all subjects in this study, we identified dense clusters of SVs with overlapping breakpoints. Most such instances could be resolved into one of four complex SV classes: tandem-duplications with nested deletions, non-tandem duplications, deletion-inversion-deletion events, and duplication-inversion-duplication events [4] (**Figure S3; Table S3**). Non-tandem duplications were the most common form of complex SV (**Table S3**), and these have not been documented in previous genome-wide studies. Insertions occurred in direct or inverted orientation with equal probability, and 22% were interchromosomal (see arrows, **Figure 1; Figure S4**). The majority (73%) had target-site deletions at the insertion point. We detect an average of 251 complex SVs per individual, thus complex SVs represent common form of genetic variation in humans.

### SV Genotyping and *de novo* Mutation Detection

Previous studies by our group and others found that *de novo* SVs occur at significantly higher rates in ASD compared to typically developing offspring [6, 3]. The more comprehensive SV dataset here provides an opportunity to investigate *de novo* structural mutation with much greater sensitivity. Identification of *de novo* SVs from WGS data, however, is a significant challenge. Given the expected number of false-positives in our call set (≥200/subject), the overwhelming majority putative de novo mutations will be errors [8]. To address this challenge, joint genotyping of SVs was performed across all samples using gtCNV, a Support Vector Machine (SVM) based algorithm we developed here to estimate genotype likelihoods for deletions and duplications (https://github.com/dantaki/gtCNV), based on multiple features including read depth, discordant paired-ends, and split-reads. Inversions deletions and insertions called by Lumpy were genotyped using SVtyper, which performs Bayesian Likelihood estimation based on discordant paired-end and split-read observations at each junction [14]. Putative *de novo* SVs were validated by microarray analysis or through PCR and Sanger sequencing (**Table S4**). We detected 31 *de novo* SVs and validated 19 in 97 offspring. *De novo* SVs consisted of a diversity of mutation classes including deletions (*n* = 11), duplications (*n* = 2), inversions (*n* = 1), *Alu* insertions (*n* = 3), and complex SVs (*n* = 2; **Table 1**), their positions are indicated by circles in the Circos plot in **Figure 1**. The overall FDR for *de novo* SV calls was 39% (12/31). This represents a substantial improvement in *de novo* SV calling accuracy, compared to the 93% FDR from a recent study by the Genome of the Netherlands (GoNL) consortium [8]. Furthermore, 12 false positive *de novo* mutations in a call set of 29,719 SV sites represents a very low error rate overall (0.04%).

### High Rate of *de novo* Structural Mutation in Humans

*De novo* SVs were observed in 19.7% of cases (95% confidence interval (CI) = 0.113-0.322) and 19.2% of controls in this study (95% CI = 7.3-42.2%), a 3-fold and 10-fold higher rate respectively than what was reported in previous studies of ASD (**Figure 2**). The higher rate of *de novo* SV is driven by increased sensitivity of our methods for detecting copy neutral and smaller SVs. The majority of de novo SVs (58%, 11/19) were undetectable using a high density (2.5 million) microarray (**Figure S5**). In contrast to previous studies, the rate of *de novo* SVs was not elevated in cases (**Figure 1**) compared to the controls in this study (*P* = 0.39) or compared to a combined set of 276 control trios from this study and from GoNL (*P* = 0.17). While the mutation rate was not elevated in cases, *de novo* SVs were larger (median length ASD = 10.9kb; controls = 1.2kb; permutation *P* = 0.026) and a greater proportion of SVs intersected an exon of at least one gene (cases = 11.1%; combined controls = 2.8%; permutation *P* = 0.01).

**Figure 2.**
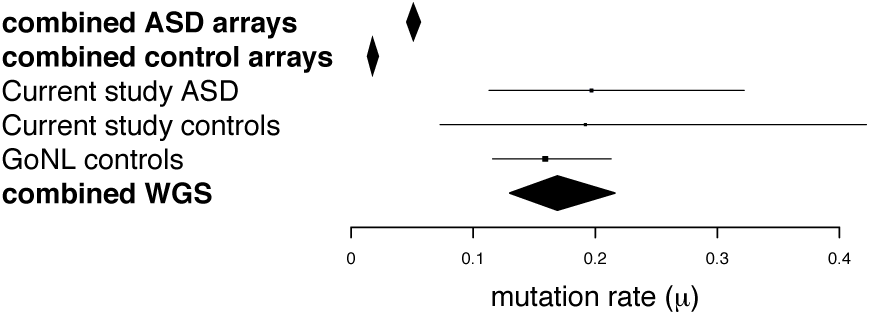
De novo Structural Mutation Frequency. Forest plot indicating the average mutation frequency per genome (*μ*). Error bars represent the 95% confidence intervals assuming a Poisson distribution, and boxes are proportional to the sample size tested.

### Fine Scale Characterization of *de novo* Structural Variants

Multilayered genetic information extracted from the genome sequences of patients can provide further insights into the origin, mutational mechanism, and functional impact of *de novo* SVs. For *de novo* events we assessed the parent of origin and the junction sequences obtained by local *de novo* assembly of breakpoints. An illustrative example, a *de novo* deletion of *CACNG2* (Calcium channel, voltage-dependent, gamma subunit 2) detected in a patient, is presented in **Figure 3**. Following detection of the deletion (Figure 3A) and genotype likelihood estimation of family members (Figure 3B), the paternal origin of the deletion could be inferred from allelic ratios of SNPs within the deleted region (Figure 3C). The complete sequence of the breakpoint junction could be assembled from reads that partially mapped near the deletion boundaries (Figure 3D). From the assembled breakpoint sequence we inferred that the deletion eliminates exon 2 and all but 634bp of intron 1 (Figure 3E) and that the deletion occurred by a non-homologous end-joining mechanism [15]. The mutant transcript lacking exon 2 of *CACNG2* was confirmed in a fibroblast line derived from the patient and is predicted to result in the inframe deletion of 30 amino acids of its extracellular AMPA receptor-binding domain (Figure 3F).

**Figure 3.**
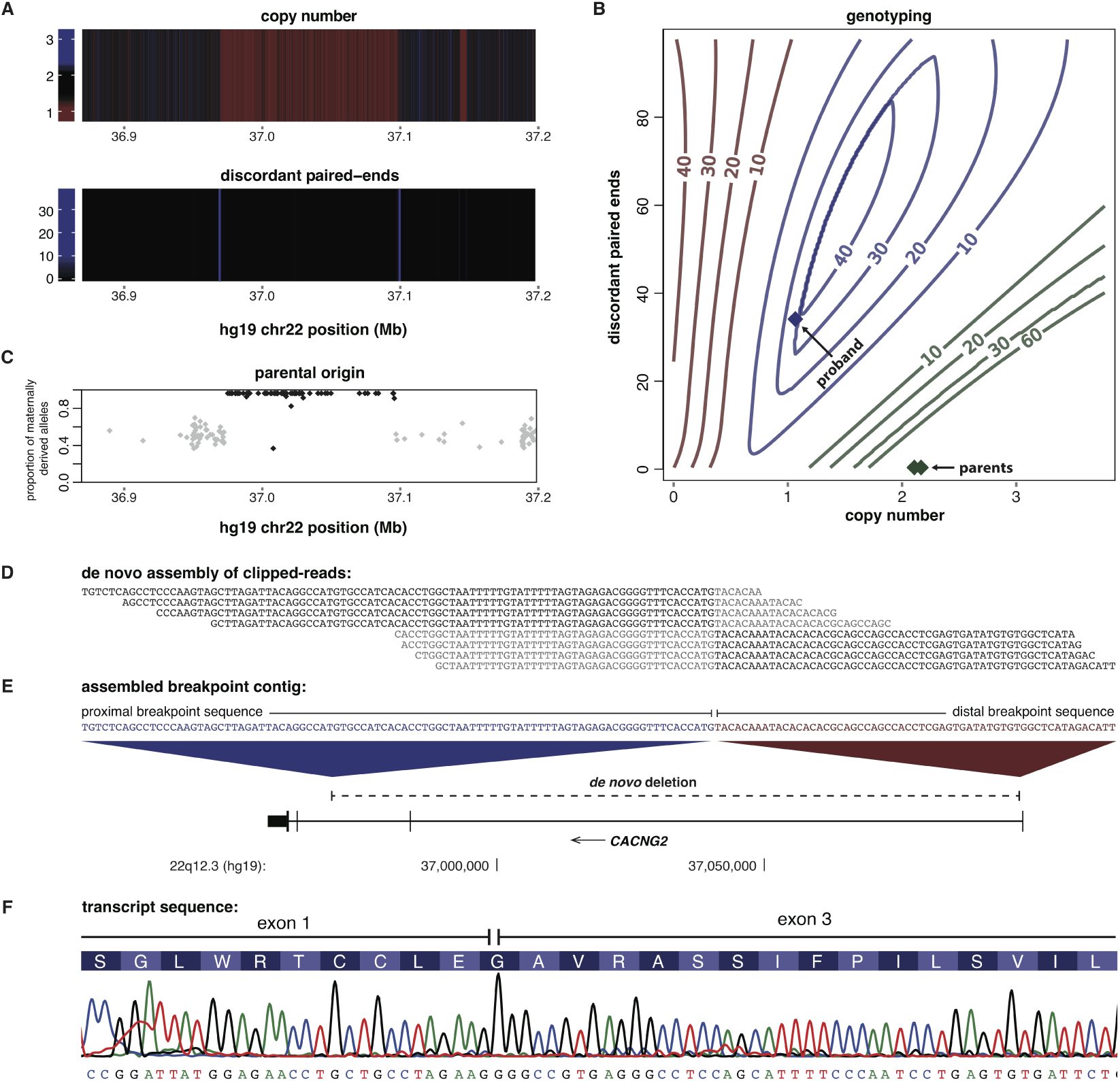
Detection, Genotyping and Sequence Characterization of de novo SVs. A) Heatmaps show a deletion signal from the total sequence coverage (copy number) and coverage of discordant paired ends. B) SVs are genotyped using a support vector machine (SVM) algorithm we developed called gtCNV. The contour plot shows the phred-scaled genotype likelihoods for homozygous reference (green), heterozygous (blue) and homozygous (red) genotypes (for simplicity only read depth and discordant paired-end features are plotted). The colored diamonds indicate the genotype likelihoods for the proband and the parents. C) A majority of SNP alleles between the deletion boundaries derive from the mother (shown in black), confirming a deletion of the paternal haplotype. D) *de novo* assembly of clipped-reads resolved the breakpoint to single base pair resolution. Unaligned sequences within clipped reads are highlighted in gray. E) Aligning the assembled contig to the genome reveals the deletion breakpoint. Unique sequence proximal and distal to the breakpoint suggests a non-homologous end joining (NHEJ) mechanism. F) The mutant transcript of *CACNG2* was sequenced from a fibroblast line derived from the patient and results in an in-frame deletion of exon 2.

**Table 1.**
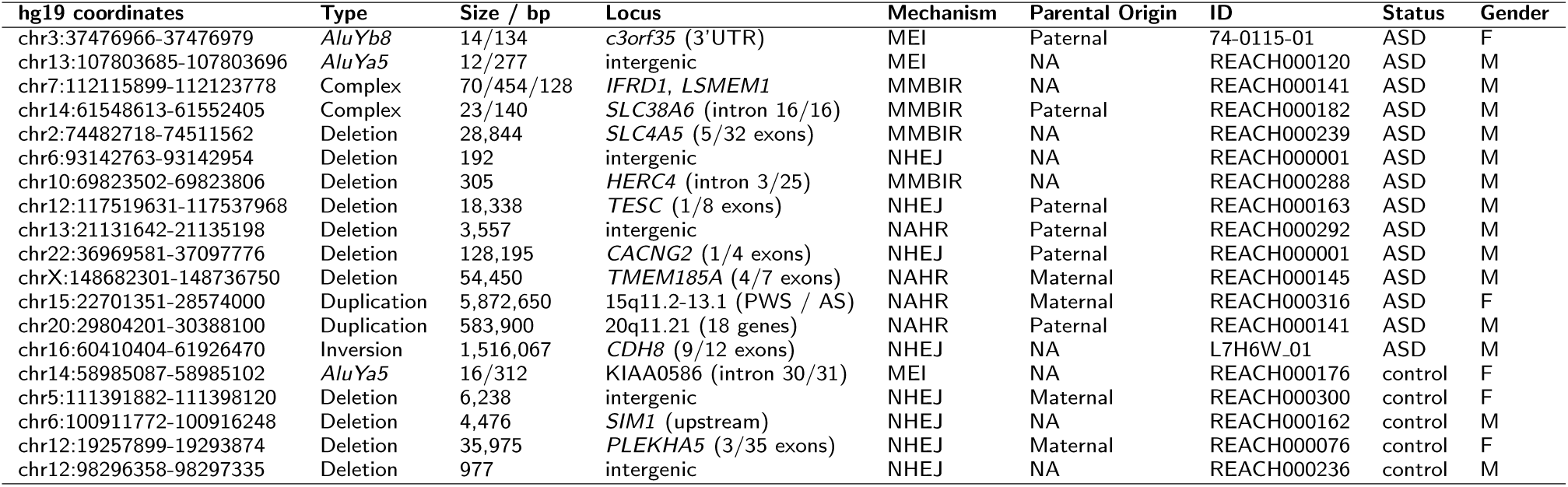
*De novo* Structural Variants.

MEIs, balanced SVs and complex rearrangements have not been systematically ascertained genome-wide in previous studies of ASD. Our results highlight how detection of these SV classes is useful for gene identification. For example, one validated MEI was a partial *AluYb8* insertion in the 3’UTR of *c3orf35* (**Figure 4**) This single observation was surprising given the low rate of *de novo* MEIs and the strong depletion of MEIs within 3’UTRs in our call set (OR = 0.44; 95% confidence interval = 0.34-0.60; **Figure S6**). Additionally, a *de novo* inversion (1.52Mb) was identified in this study that disrupts cadherin-8 (*CDH8*; **Figure 4**). These results strengthen the evidence from previous studies implicating *c3orf35* [6] and cadherin-8 [16] in ASD.

**Figure 4.**
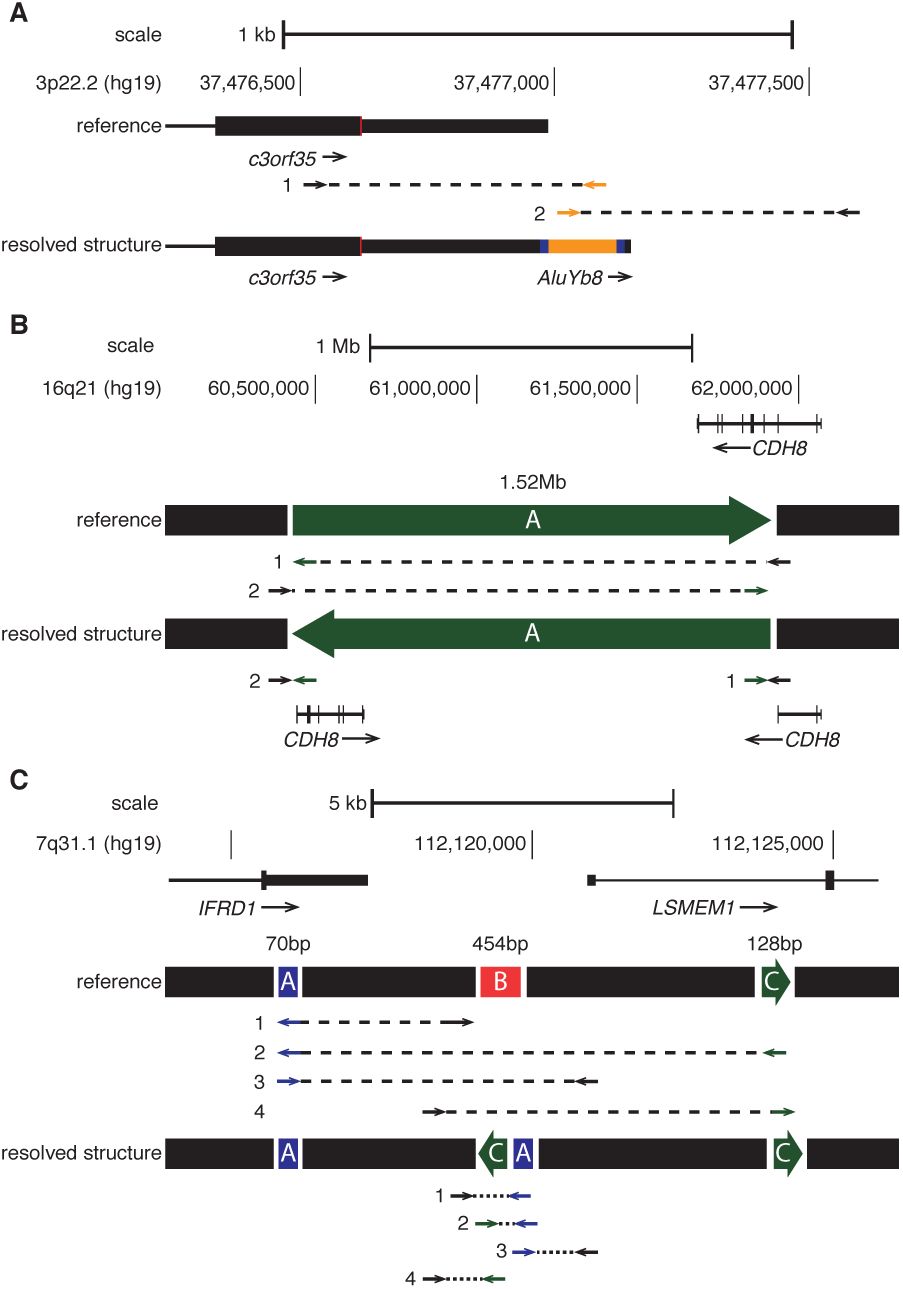
De novo Structural Mutations of Genes Detectable Through Genome Sequencing. Discordant paired-end mapping identifies *de novo* structural variants. A) *AluYb8* insertion into the 3’ UTR of *c3orf35* (chromosome 3 open reading frame 35), with a 14bp target site duplication (shown in blue). B) A simple inversion with a distal breakpoint in intron 3 of *CDH8.* C) A complex SV at the promoter of *LSMEM1* (n.b. segments not shown to scale). Arrows indicate the discordant orientation and location of paired-end reads relative to the hg19 reference genome and the concordant pattern of paired end reads relative to the resolved structure. Black segments are unchanged in the SV events, green segments are inverted, blue segments are duplicated, and red segments are deleted.

### Complex Mutation Clusters

The complexity of *de novo* SVs consisted not only of clusters of deletions, duplications and inversions occurring as single events (**Figure 4**), but also included co-occuring *de novo* nucleotide substitutions and indels in the surrounding region (**Figure 5**). We observed greater clustering of *de novo* SVs and point mutations within individual genomes than would be expected by chance. In total, six *de novo* mutations (five SNVs and one indel) were located within 100kb of *de novo* SV breakpoints in this study, a 63-fold enrichment compared to random mutation (permutation *P* = 0.00053; **Table S5**). Adjacent *de novo* SVs and SNVs were located tens of kilobases apart; therefore, the enrichment of *de novo* substitutions around SV breakpoints could not be explained as an artifact due to the mismapping of reads at the junction. **Figure 5** illustrates examples of complex mutation clusters identified in two individuals.

**Figure 5.**
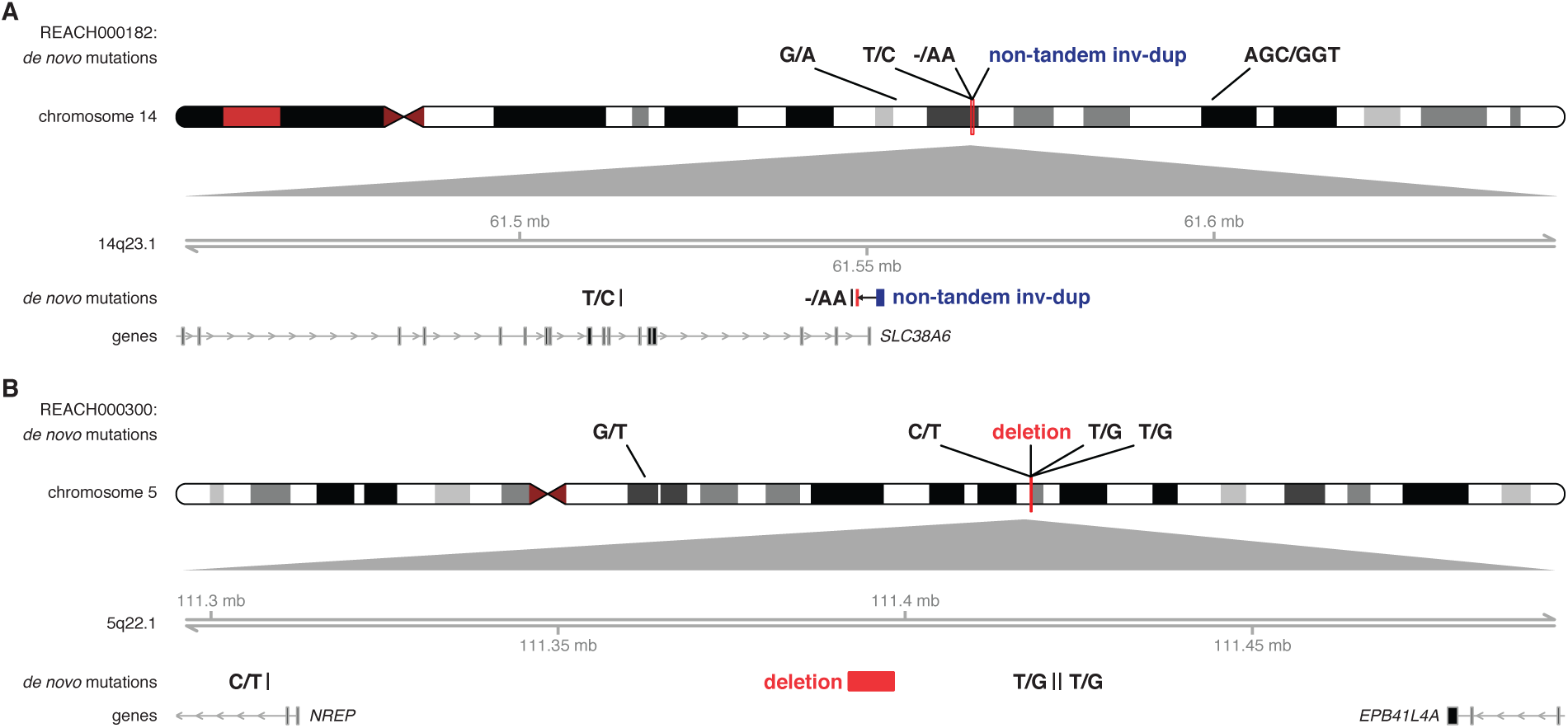
Mutational Clustering of SVs, Indels, and SNVs. Two examples of complex mutation clusters are shown in individuals. A) REACH000182, a 143bp sequence near the 3’UTR of *SLC38A6* that was duplicated and inserted into the first intron of the gene with a concomitant deletion of 23bp at the insertion site. Additionally, a *de novo* indel and SNV occurred at 211bp and 34,611bp proximal to the insertion site, B) REACH000300, a 6.2kb *de novo* deletion was detected at 5q22.2 and three *de novo* SVs occurred within a 100kb of the breakpoints. The 200kb zoomed in locus below the ideogram shows the positions of *de novo* mutations relative to each other. Gene tracks below the mutation show the longest transcript of each gene within the locus, with arrows indicating the strand and bars indicating the exons of genes.

### Pathogenic Inherited Structural Variants

We examined the call set for known pathogenic SVs and observed five rare or *de novo* CNVs in cases overlapping with known ASD or intellectual disability (ID) risk variants (expected = 2; 95 CI = 0-5; *P* = 0.063; OR = 2.42). We did not observe significant overlap with genes impacted by *de novo* loss of function variants identified in ASD and ID by exome sequencing (observed = 13; expected = 14.9; 95 CI = 8-23; *P* = 0.72; OR = 0.84). Inherited risk variants were identified in four unrelated cases (6%). One *de novo* SV identified in this study, a duplication of 15q11.2-13.1 (**Table 1**) has also been previously implicated in ASD [17]. We observed two paternally inherited deletions of 15q11.2 (that include *CYFIP1*) [18]. Inherited X-linked variants include a duplication-inversion-duplication event at Xp21.1-21.2 that duplicates the Dp71 isoform of *DMD* (dystrophin) and disrupts *TAB3* (**Figure S3**). Duplications and deletions of *DMD* are associated with Duchenne muscular dystrophy, and some alleles may predispose to ASD [18, 19, 20]. Lastly, we detected a maternally inherited deletion of *NRXN1* (neurexin 1) [18]. Follow up genetic analysis of the extended family revealed that the deletion occurred *de novo* in the mother (**Figure S7**) and was not carried by the maternal grandparents. This observation highlights that, while these disease-associated variants are inherited from a parent, the SVs described above occur within regions that are prone to frequent recurrent rearrangements and are likely to be mutations that occurred in recent ancestry.

## Discussion

We have assembled the most complete set of SVs in ASD to date. Whole Genome Sequencing of trio families reveals a diverse landscape of structural variation throughout the genome and reveals a higher rate and complexity of structural mutation than previously recognized. Structural mutations detected in patients include previously undetectable events that disrupt genes and are likely to influence disease risk.

The combined frequency of *de novo* SVs that we observe (*μ* = 0.195) is more than triple the estimate from a previous WGS study of autism (*P* = 0.058) [9]. Our estimate is also slightly higher than the rate observed in a family based study by the GoNL consortium (*μ* = 0.16) [8]. The mutation rate reported here will ultimately prove to be an underestimate as well due to the challenges of detecting SVs using short read next-generation sequencing technology.

With the improved ascertainment of small deletions, inversions and MEIs, we observe a similar overall mutation rate in cases and controls, in contrast to previous studies by our group [3] and others that were based on microarray technology [21, 6]. Thus, a genetic contribution of *de novo* SV to ASD is evident, not from an elevated frequency of genomic rearrangement, but instead from the greater proportion of new mutations that disrupt genes. In this respect, the contribution of *de novo* structural mutation to ASD bears similarity to that of *de novo* loss-of-function mutations detected by exome sequencing [22, 23].

Studies of genetic diversity in populations reveal a diverse spectrum of SV [7], but do not fully illuminate the mutational process that gives rise to that diversity. Here we show that one third of *de novo* SVs consist of mobile elements, balanced, or complex mutations, underlining the role that these mutational mechanisms play in generating genetic diversity and disease risk. Candidate loci for ASD were identified from two such *de novo* SVs, including a mobile element insertion in *c3orf35* and an inversion disrupting cadherin-8. Published studies of ASD provide additional evidence implicating both genes, including a *de novo* deletion disrupting *c3orf35* [5] and segregating deletions of *CDH8* observed in ASD families [16].

Our results highlight how clusters of SVs arise through complex mutation events that generate combinations of deletions, duplications, insertions and inversions (and sometimes all of the above). Adding further complexity to the mutational process, we show that 16% (3/19) of *de novo* SVs co-occur with clusters of *de novo* SNVs and indels. These results expand upon our previous study reporting the observation of *de novo* nucleotide substitution ’showers’ [24]. Our current findings suggest that sequence variation and structural variation can arise from common mechanisms. We hypothesize that such complex mutation clusters form as a consequence of break-induced replication (BIR) during double-stranded break (DSB) repair. BIR is significantly more error prone than normal DNA replication and occurs over hundreds of kilobases [25], a scale that is similar to the length of the mutation clusters observed here.

The observation of complex mutation clusters is interesting in light of a previous study from the 1KG project, which found that SNPs and indels in the population are enriched within 400kb of deletion breakpoints. It was further hypothesized that the observed enrichment of SNPs and indels is due to relaxed selection at these loci [26]. Based on our results, we suggest that the observed correlation of SNPs and SVs is in part attributable to the underlying mutational processes and is not driven entirely by selection.

With our high coverage and complementary SV discovery methods, we are able to detect 5,213 SVs per individual, 27% more alleles per genome than we and others recently reported in the 1KG call set (*n* = 4,095 / genome) [7]. However, neither study presents a complete catalogue of SVs. For example, the substantial non-overlap of duplications between call sets in this study and the 1KG project suggests that both have underestimated the number of duplications. The short-read shotgun sequencing technology used here still possesses significant technical limitations that impede our ascertainment of SVs. Application of new long-read sequencing technologies [27], will be another significant step towards uncovering the complexity of structural variation in autism.

## Methods

### Patient Recruitment

Patients were primarily referred from clinical departments at Rady Children’s Hospital, including the Autism Discovery Institute, Psychiatry, Neurology, Speech and Occupational Therapy and the Developmental Evaluation Clinic (DEC). Further referrals came directly through our project website (http://reachproject.ucsd.edu). The Autism Center of Excellence at the University of California San Diego contributed a further 11 trios (http://www.autism-center.ucsd.edu). Each child included in the study has an existing ASD diagnosis, and has been comprehensively evaluated by licensed clinicians utilizing a battery of tests such as the Autism Diagnostic Observation Schedule (ADOS) [28]. Prior to appointments, families were provided with Institutional Review Board (IRB) approved consent forms and Health Insurance Portability and Accountability (HIPAA) consents. DNA was obtained from 5mL blood draws. We recalled a subset of patients with specific genetic findings to confirm the original ASD diagnoses. These included patients with SVs in *TMEM185A, TESC, NRXN1,* and *CACNG2.* A diagnosis of ASD was confirmed in all cases.

### Whole Genome Sequencing

WGS was performed on 246 samples, which included 11 monozygotic twin pairs. One sibling from each twin pair was excluded from the dataset, which brought the final sample size to 235. WGS of 206 samples was performed at Illumina Fast Track service laboratory in San Diego using the Illumina HiSeq Platform. For 161 samples, preparations consisted of 313bp libraries and 100bp paired-end reads. For the remaining 45 samples, library size and read length were 493bp and 125bp respectively. In addition, a subset of our data consisted of 40 samples sequenced at the Beijing Genomics Institute (BGI) using the Illumina HiSeq as described previously (SVs were not reported in this publication) [24], and genomes were realigned using Burrows-Wheeler Aligner (BWA-mem version 0.7.12) to the hg19 reference genome [29].

Sequence alignment and variant calls were generated on families using our WGS analysis pipeline implemented on the Triton compute cluster at UCSD (http://tritonresource.sdsc.edu/). Short reads were mapped to the hg19 reference genome by BWA-mem version 0.7.12 [29]. Subsequent processing was carried out using SAMtools version 1.2 [30], GATK version 3.3 [31], and Picard tools version 1.129 (http://broadinstittute.github.io/picard), which consisted of the following steps: sorting and merging of the BAM files, indel realignment, fixing mate pairs, removal of duplicate reads, base quality score recalibration for each individual [32].

### SV Detection

We utilized three complementary algorithms to detect SVs. ForestSV is a statistical learning approach that integrates multiple features from WGS data, including read depth and discordant paired-end signal, to identify deletions and duplications [11]. Lumpy uses discordant paired-end and split-read signal to identify breakpoints for deletions, duplications, inversions, translocations, and complex SVs [14, 12]. Finally, Mobster uses discordant paired-end and split-read signal in combination with consensus sequences of known active transpsosable elements to identify mobile element insertions (MEIs) [13].

### SV Post-Processing

We assembled call sets of deletions, duplications, inversions, complex structural variants, and mobile element insertions detected in 246 individuals. For monozygotic twins we generated a consensus call set for each twin pair from the raw SV calls as an initial processing step.

### ForestSV

Initial QC were applied to the raw ForestSV including removal of calls that overlapped by 50% with centromeres, segmental duplications, regions with low mappability, and regions subject to somatic V(D)J recombination (T-cell receptors and immunoglobulin genes). Large SVs that were fragmented into multiple calls by ForestSV were stitched together as a function of their separation distance divided by the total length of the individual calls. SVs between individuals were collapsed based on ≥50% reciprocal overlap and the same median start and end coordinates were assigned to each call.

### Lumpy

Structural variants were called within families using the Speedseq SV pipeline (v0.0.3a), which processes the samples using Lumpy (v0.2.9), and genotypes variants using SVtyper (v0.0.2) [14, 12]. The pipeline outputs deletions, duplications, inversions, and breakpoints that cannot be assigned to one of the three classes. To detect complex SVs, we wrote a custom algorithm to cluster overlapping pairs of breakpoints and resolve the patterns and ordering of breakpoint alignments to the reference genome. Calls between individuals were considered to be the same SV if they shared the same start and end coordinates within a margin of error defined by the read length (100bp for most samples). SVs were filtered if one or more breakpoint overlapped regions with low mappability, including centromeres and segmental duplications, and also regions subject to somatic V(D)J recombination (T-cell receptors and immunoglobulin genes).

### Mobster

Mobile elements were called by Mobster within families and were included in the call set if they had at least five reads supporting the call in one individual in the family, including discordant paired-ends at both the 3’ and 5’ sides of the insert point [13]. Calls between individuals were considered to be the same SV if they shared the same insertion coordinates within a margin of error defined by the read length (100bp for most samples). SVs were filtered if one or more breakpoint overlapped regions with low mappability, including centromeres and segmental duplications, and also regions subject to somatic V(D)J recombination (T-cell receptors and immunoglobulin genes).

### SV Genotyping and *de novo* Mutation Calling

We utilized two methods for assigning genotype likelihoods for structural variants, gtCNV and SVtyper [14], which are detailed below. We then derive a quality score for each SV site, defined as the median genotype likelihood for individuals genotyped as non-reference.

The finalized call sets were filtered solely on quality scores, using thresholds of 12 for deletions genotyped with gtCNV, 8 for duplications genotyped with gtCNV, and 100 for SV breakpoints genotyped with SVtyper. The FDR of the combined call set was estimated from Illumina 2.5M SNP array data using the Intensity Rank Sum test implemented using the Structural Variation Toolkit (http://svtoolkit.sourceforge.net/). From the finalized call set we extracted *de novo* mutations that had a non-reference genotype in the child and reference genotypes in both parents, with a parent allele frequency of zero in the cohort.

### gtCNV

We developed a likelihood-based method to classify CNV genotypes, gtCNV, a Support Vector Machine (SVM) machine-learning approach that genotypes deletions and duplications. The classifier was trained on high coverage CNV data from nine trios sequenced as part of the 1KG project.

We selected read depth, discordant paired-ends, and split-reads as features when training the SVM. Features were extracted for all deletion and duplication calls made by ForestSV and Lumpy. When determining coverage, we masked regions overlapping with segmental duplications. For each SV, we calculated mean coverage, which was then normalized to the mean chromosomal coverage for each sample. We also extracted all discordant paired-ends and split reads (mapping quality ≥20) using the samtools API for Python, implemented in pysam (https://github.com/pysam-developers/pysam [30]). Discordant paired-ends were defined as reads with insert sizes more than five standard deviations from the mean.

The SVM training utilized a Radial Basis Function (RBF) kernel, implemented in Python using scikit-learn [33, 34]. In order to determine the optimal RBF kernel parameters, we estimated the false-discovery rate (FDR) for deletions and duplications in the call set using the Intensity Rank Sum (IRS) test. Optimal parameters were C = 1 and gamma = 0.005, which had an FDR of 5.9% for deletions at a quality score of ≥12. The optimum parameters for duplications were C = 1 and gamma = 0.01, which had an FDR of 6.0% at a quality score of ≥8. The gtCNV software can be found on github (https://github.com/dantaki/gtCNV) and the method will be further detailed in a companion paper to follow in the near future.

### SVtyper

Genotyping of Lumpy calls was performed using SV-typer, implemented as part of the speedseq SV pipeline [14]. SVtyper is a Bayesian SV breakpoint genotyping algorithm that estimates the likelihood that a genotype is non-reference (SQ) based on allele counts at each junction. A quality score for each individual SV locus was derived based on the median genotype likelihood for individuals genotyped as non-reference. An optimal quality score threshold for Lumpy was determined as described in the section above. We performed family-based calling and genotyping for Lumpy calls, and kept variants that had a median quality score ≥100 across the cohort. For complex variants with multiple overlapping breakpoints, we kept variants that had a median quality score ≥100 for at least one breakpoint.

The performance of SVtyper was assessed using intensity rank sum test described above. The FDR of CNV sites was 3.3% for deletions, 9.5% for tandem duplications, 0% for deletions in complex events, and 11.5% for duplications in complex events (complex combined FDR = 7.5%).

### Detection of *de novo* SNVs and Indels

Putative *de novo* SNVs were called using ForestDNM, a custom machine-learning pipeline that uses a random forest classifier to predict the validation status of putative *de novo* SNVs identified by GATK Unified Genotyper [24]. Putative *de novo* indels were called using three different algorithms, GATK, Platypus, and Scalpel [31, 35, 36]. We called variants first genome-wide using GATK and Platypus. Then we used Scalpel for targeted *de novo* assembly of the locus around this set of putative *de novo* indels. We kept *de novo* indels called by at least two out of three algorithms. We then excluded any indels observed in the GATK or Platypus variant call files (VCF) of the entire cohort more than once, or indels that are common in the population from 1000 Genomes Project data. The genome-wide burden of *de novo* SNVs in cases and controls was 66.9 and 63.3 respectively, for indels it was 6.67 and 6.11. Analysis of *de novo* SNVs and indels will feature in a future publication.

### SNP Microarray CNV Validation

We performed genome-wide assessment for copy number variation (CNVs) in the majority of individuals in this study (*n* = 205) via Illumina 2.5 million singlenucleotide polymorphism (SNP) microarrays. CNVs were detected using trio-based calling implemented in the PennCNV algorithm [37], and retained if they had at least eight supporting probes.

### PCR Validation Structural Mutations

We designed PCR primers flanking breakpoints for complex and balanced structural variants (*n* = 3), all of which we successfully validated. We attempted validation of nine putative *de novo* MEIs, six *Alu* and 3 L1 insertions. Primers were designed flanking the insertion point for *Alu* elements, and for L1 elements one primer was designed flanking the insertion point, and two within the element (both sense and antisense since the orientations of the insertions were unknown). PCR amplification validated three *de novo Alu* elements when run on an agarose gel; the remaining putative *de novo* variants were false positives. PCR products were cloned using TOPO-TA vectors. Resulting clones were screened and sequenced using M13 primers from both ends of the vector insert. The subfamily was assigned by comparing the sequence results with the consensus *Alu* sequences using BLAST.

### Assembly of Breakpoints

For deletion and duplication SVs, we performed *de novo* assembly of clipped-reads using Velvet [38], and determined the precise breakpoint down to a single base pair resolution for 60.8% of deletions (*n* = 11,168), and observed that 17.9% of deletions have inserted sequence at the breakpoint. For duplications we determined the breakpoints for 31% (*n* = 733). Breakpoint positions were assigned to SV coordinates where applicable in **Supplementary Data Set 1**.

### SV Burden

The burden of *de novo* structural variants between individuals with ASD in this study and the combined controls from this study and GoNL was assessed using a case-control permutation test implemented in PLINK [39].

### MEI Permutations

To permute the enrichment/depletion of MEI insertions in genomic features, we shuffled the position of the observed MEI sites across the genome using BedTools [40], maintaining the orientation of the MEI (sense or antisense), while excluding any overlap with sequencing gaps. We counted the number of times where a shuffled MEI overlaps the following genomic features: exons, introns (sense and antisense orientation separately), promoters / 5’UTRs, and 3’UTRs. We performed 10,000 permutations and compared the observed overlap to the expected overlap.

### Overlap of SVs with Known Polymorphic SV Events

Deletions, duplications, and inversions were intersected with the 1KG call set using BedTools, and considered part of the same polymorphic or recurrent SV event if they had ≥50% reciprocal overlap. MEIs were considered to overlap if their insertion point was located within 100bp of an MEI event of the same class from the 1KG integrated SV set or the database of retrotransposon insertion polymorphisms.

### Overlap of SVs with Published CNV Data

We permuted the expected overlap of SVs with CNV regions (CNVRs) previously associated with ASD, intellectual disability, and developmental delay, derived from two large scale microarray CNV studies [41, 21]. These CNVRs are significantly enriched in cases versus controls, and are either hotspots flanked by segmental duplications or peaks of enrichment derived from the intersection of multiple breakpoints. We shuffled the position of the observed rare SVs in children (including SVs that are *de novo* or have a frequency ≤1% in parents) randomly using BedTools, maintaining the size of the CNVs and the chromosome, while excluding any overlap with sequencing gaps. We counted the number of times where at least 90% of a shuffled CNV overlaps a CNVR. When a single gene has been implicated by a CNVR, we stipulated that the CNV has to overlap only one exon of the gene to be counted. This method is conservative because it allows for small CNVs overlapping only a small proportion of larger CNVRs to be counted, i.e. we do not require the overlap to be reciprocal.

When performing gene-set enrichment analysis with published exome sequencing data, we determined the number of SVs overlapping with genes impacted by one or more loss of function SNVs and indels in studies of ASD and intellectual disability, and then permuted the SV positions, maintaining the total number of genes disrupted.

### Mutational Clustering

To assess whether *de novo* SVs cluster with nucleotide substitutions or indels, we used a window based permutation approach. We took windows of 100bp, 1kb, 10kb, 100kb, 1Mb, and 10Mb around the breakpoints of *de novo* SVs and intersected the windows with *de novo* SNVs and indels in the same individuals. We then shuffled the position of these windows in the genome randomly using BedTools and calculated the expected number of window overlapping DNMs using 100,000 permutations.

### Transmission Disequilibrium Test

We tested whether variants private to families in our study and not present in the 1KG call set were transmitted to affected children more than expected by chance, using a haplotype-based group-wise transmission disequilibrium test (gTDT) [42], assuming an additive model.

## Author’s contributions

Conceptualization, J.S., W.M.B; Methodology, W.M.B., D.A., M.G., A.N., J.S.; Software, D.A., W.M.B., M.G., A.N.; Validation, G.R., T.C., D.J.B., O.H., M.B., T.S., A.G.M., A.C.W.; Formal Analysis, W.M.B, D.A., M.G., G.N.L., L.M.I., S.J.S., S.M.L.; Writing – Original Draft, J.S., W.M.B.; Writing – Review and Editing, L.M.I, S.J.S, E.C., K.P.; Resources, K.K.V., G.E.R., L.C.W., J.A.E., E.C., C.C., N.A., A.R.M., K.P., C.M.S., R.O., K.Z.; Visualization, W.M.B., D.A.; Supervision, J.S.; Project Administration, J.E., T.E.G., O.H., K.V.F.F., A.B.; Funding Acquisition, J.S.

## Acknowledgements

We would like to thank the families who volunteered for the study. Grants to JS from the NIH (MH076431, HG007497) and the Simons foundation (SFARI 275724). Grants to L.M.I. from the NIH (HD065288, MH104766 and MH105524). W.M.B. was supported by a fellowship from the Autism Science Foundation. D.A. was supported by NIH predoctoral training grant T32 GM008666. We would like to thank Wayne Pfeiffer, Mahidhar Tatineni, and the San Diego Supercomputer Center for hosting the computing infrastructure necessary to complete this project.

## Supplementary Data Sets

**Supplementary Data Set 1.** Combined call set of deletions, duplications, inversions, and complex SVs in VCF format.

**Supplementary Data Set 2.** Putative *de novo* mutations identified from the combined call set in VCF format.

### Supplementary Tables

**Table S1.** Sample Information for 235 Individuals Sequenced.

**Table S2.** Mobile Element Insertion Call Set.

**Table S3.** Descriptive Statistics for 11 Classes of Structural Variant.

**Table S4.** PCR Validation of Structural Variants.

**Table S5.** Permutations of Overlap of *De Novo* Indels and SNVs with Windows around SV Breakpoints.

**Figure S1.**
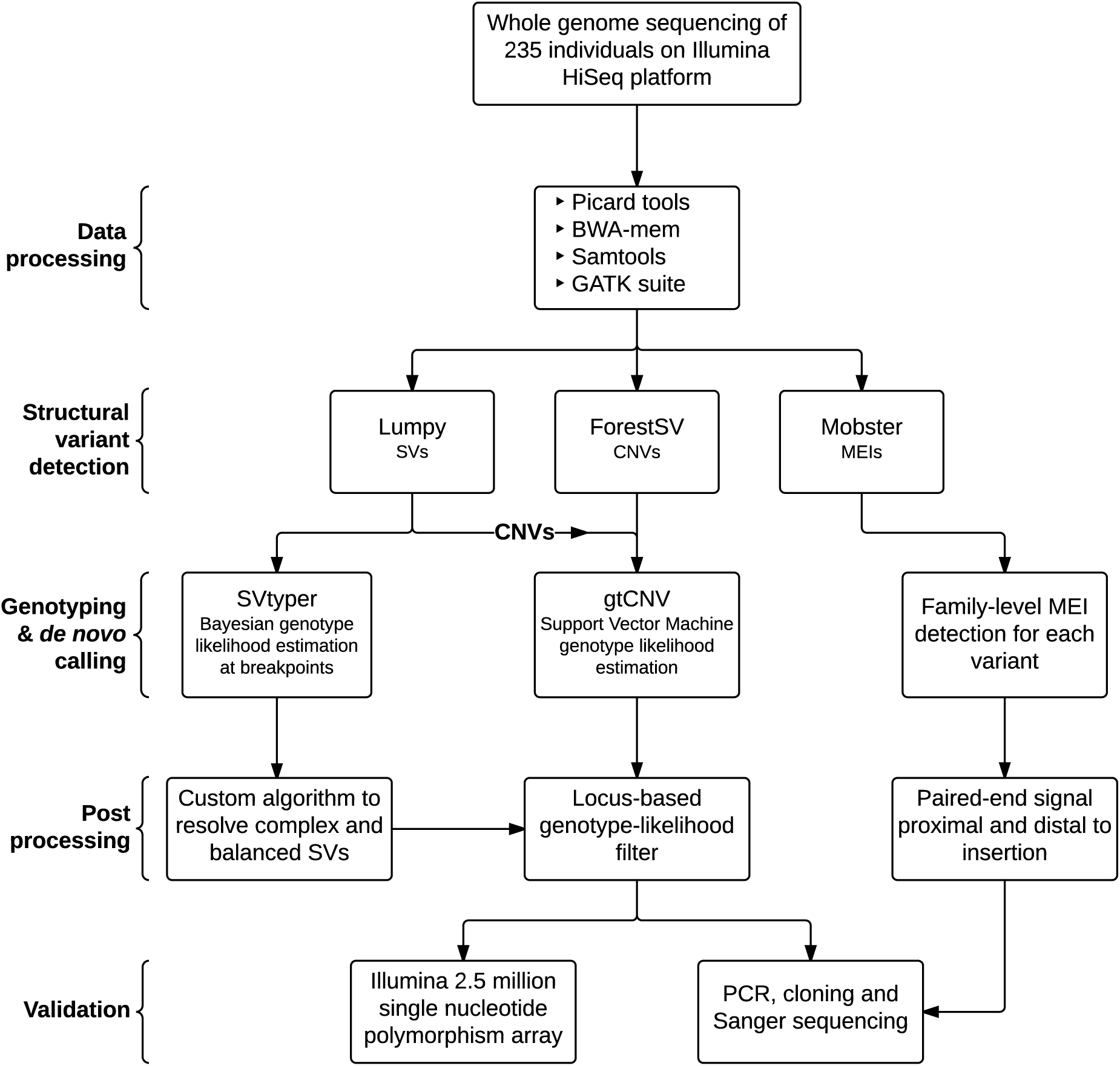
Structural Variant Discovery Pipeline. Flowchart detailing our custom pipeline for the discovery, genotyping, and validation of structural variants and de novo mutations. CNV = Copy Number Variant; SV = Structural Variant; MEI = Mobile Element Insertion; PCR = Polymerase Chain Reaction.

**Figure S2.**
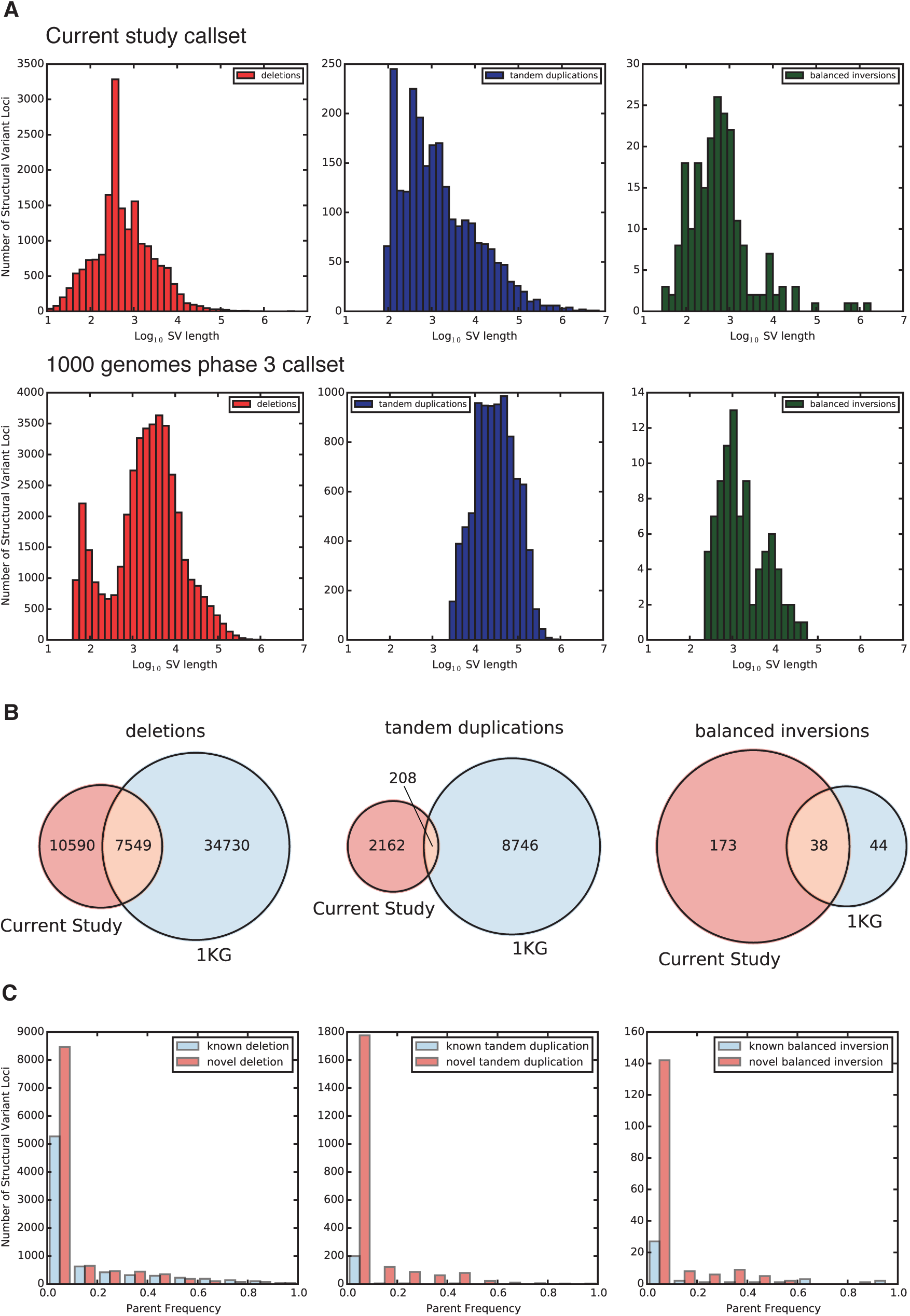
Comparison between current study call set and 1000 Genomes Phase 3. A) Histograms of the log_10_ structural variant (SV) size distributions for deletions, tandem duplications and balanced inversions in our study and 1000 genomes phase 3 (1KG) SV call set. B) Venn diagrams showing the overlap of deletions, tandem duplications and inversions detected in our study with SVs from the 1KG SV call set. C) Histograms showing the number of novel versus known SVs across a range of parent frequencies.

**Figure S3.**
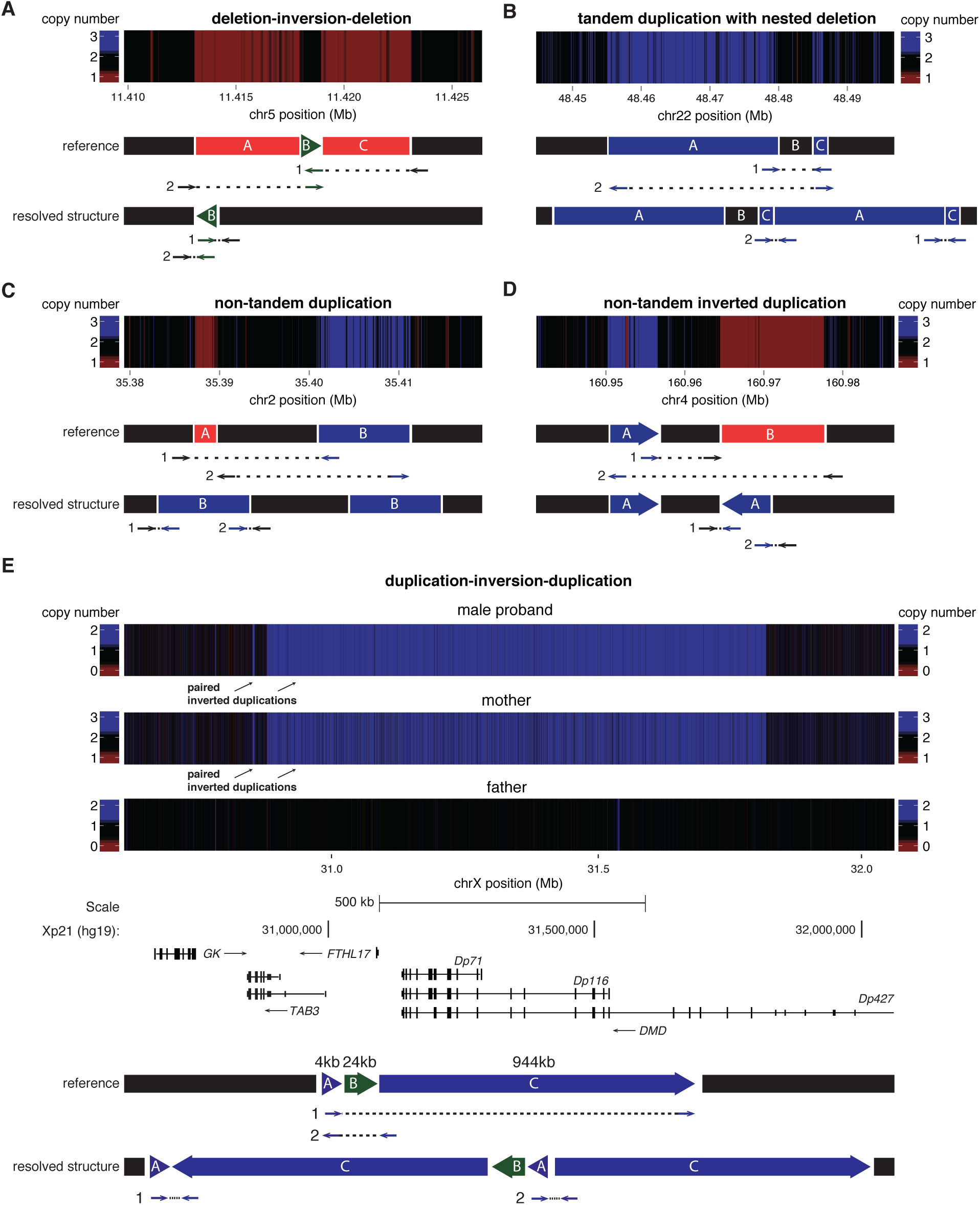
Complex Structural Variation Detected using Genome Sequencing. Examples of each class are taken from our call set of SVs. A) deletion-inversion-deletion, B) tandem duplication with nested deletion, C) non-tandem duplication, D) non-tandem inverted duplication, E) duplication-inversion-duplication (including genes in the vicinity of the SV event). Heat maps indicate changes in copy number observed from the depth of coverage at each locus, normalized to the chromosomal average. Lettered segments indicate the structure of the chromosome in the reference and the observed genome. Black segments are unchanged in the SV events, green segments are inverted, blue segments are duplicated, and red segments are deleted. Arrows indicate the discordant orientation and location of paired-end reads relative to the hg19 reference genome and the concordant pattern of paired end reads relative to the resolved structure. n.b. segments not shown to scale.

**Figure S4.**
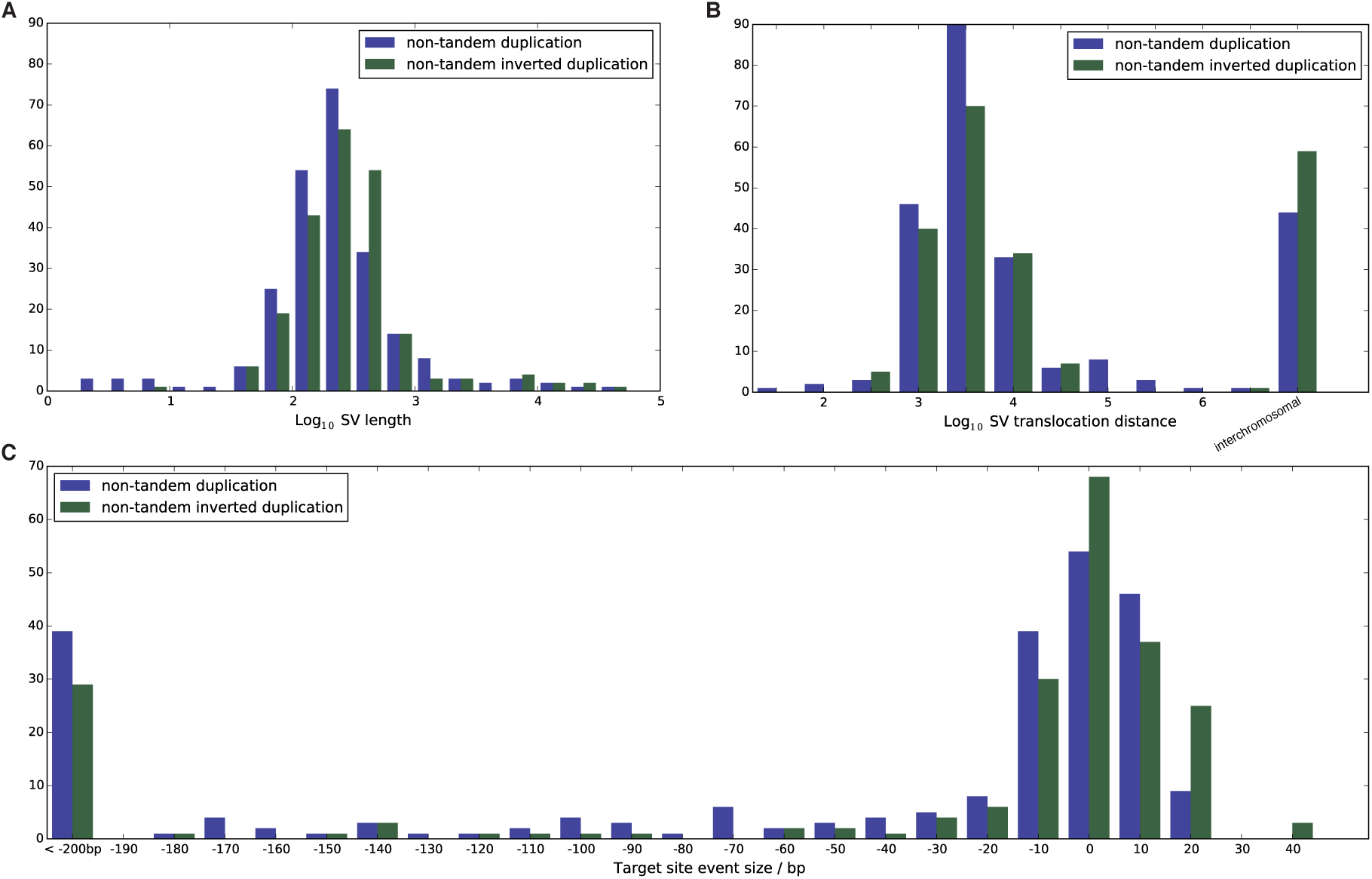
Distribution of Non-Tandem Structural Variants. a) Histogram of the lengths of non-tandem duplications (blue) and non-tandem inverted duplications (green). b) Histogram of translocation distances. c) Histogram of target site deletions or duplications at the non-tandem event’s insertion point.

**Figure S5.**
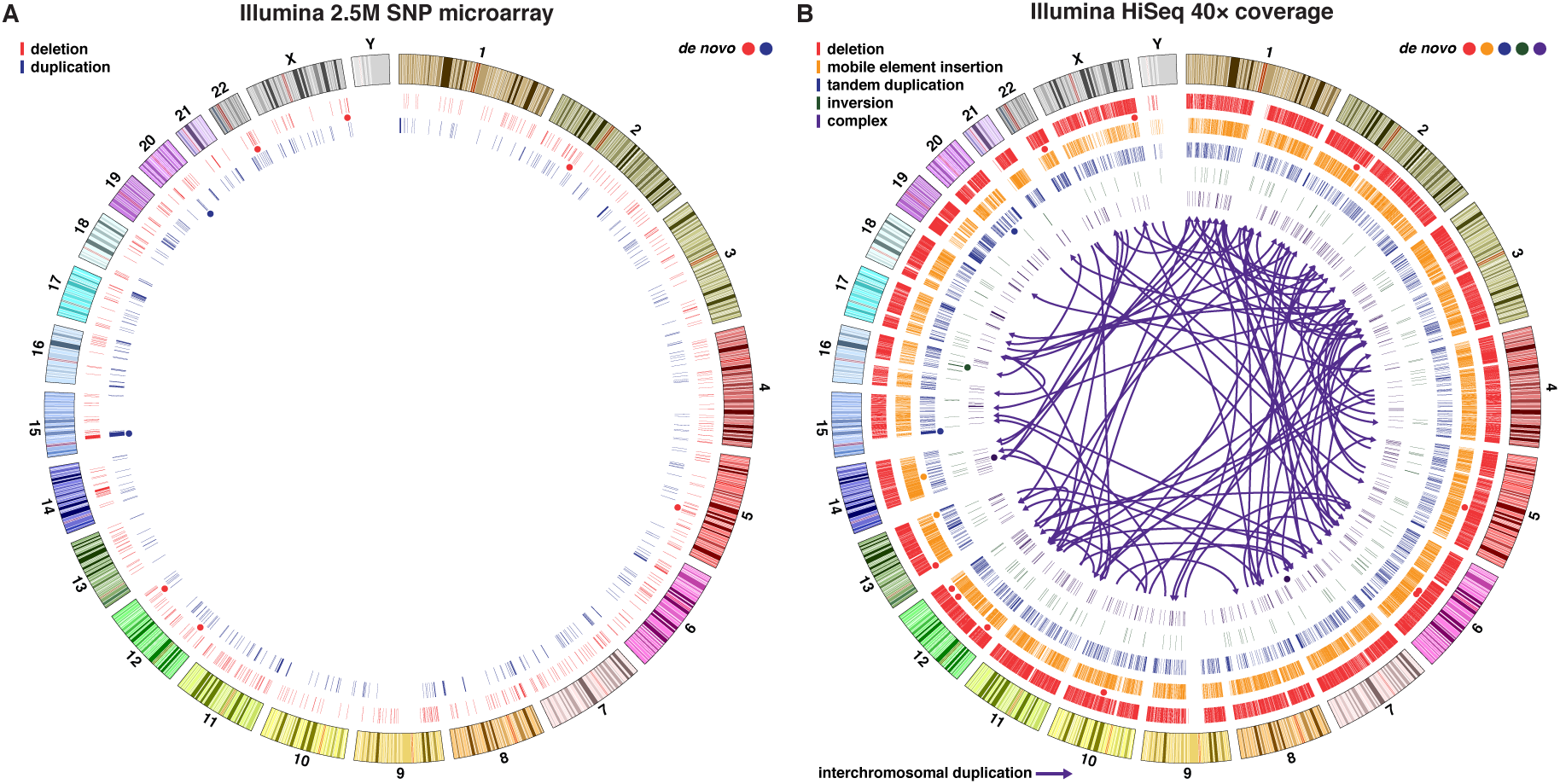
Comparison of Structural Variation Detection using Microarrays and Genome Sequencing. Circos plots comparing structural variant calls for 205 individuals in this study derived from a) Illumina 2.5 million single nucleotide polymorphism (SNP) microarray, and b) from WGS at 40× coverage on the Illumina HiSeq platform. Concentric circles represent from outermost to inner in panel: ideogram of the human genome with karyotype bands (hg19), deletions, mobile element insertions (four different classes), tandem duplications, balanced inversions, complex structural variants (four different classes). The circles indicate the location of de novo SVs, and their colors match the five SV types. Arrows represent interchromosomal duplications.

**Figure S6.**
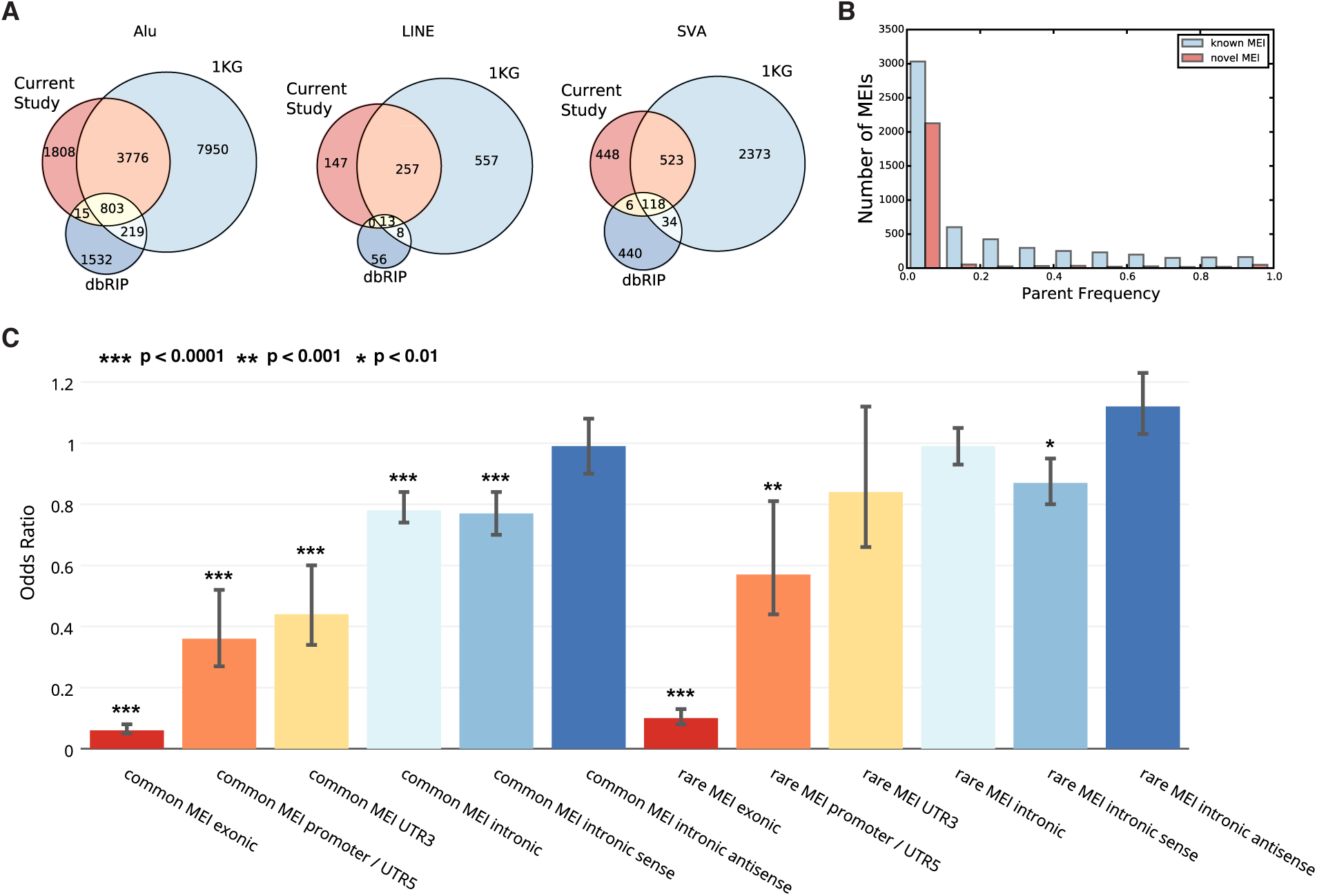
Mobile Element Insertion Overlap with Published Databases and Genomic Features. A) Venn diagrams showing the overlap of MEIs detected in our study with MEIs from the 1000 genomes project (1KG) phase 3 integrated SV call set and the database of retrotransposon insertion polymorphisms (dbRIP), calls were considered to overlap if they were within 100 base-pairs of each other. B) Histogram showing the number of novel versus known MEIs across a range of parent frequencies. C) Bar chart showing the odds ratio of the overlap of observed common (frequency ≥5%) and rare MEIs with genomic functional elements compared to expected overlap through permutation. Error bars represent the 95% confidence interval for odds ratio.

**Figure S7.**
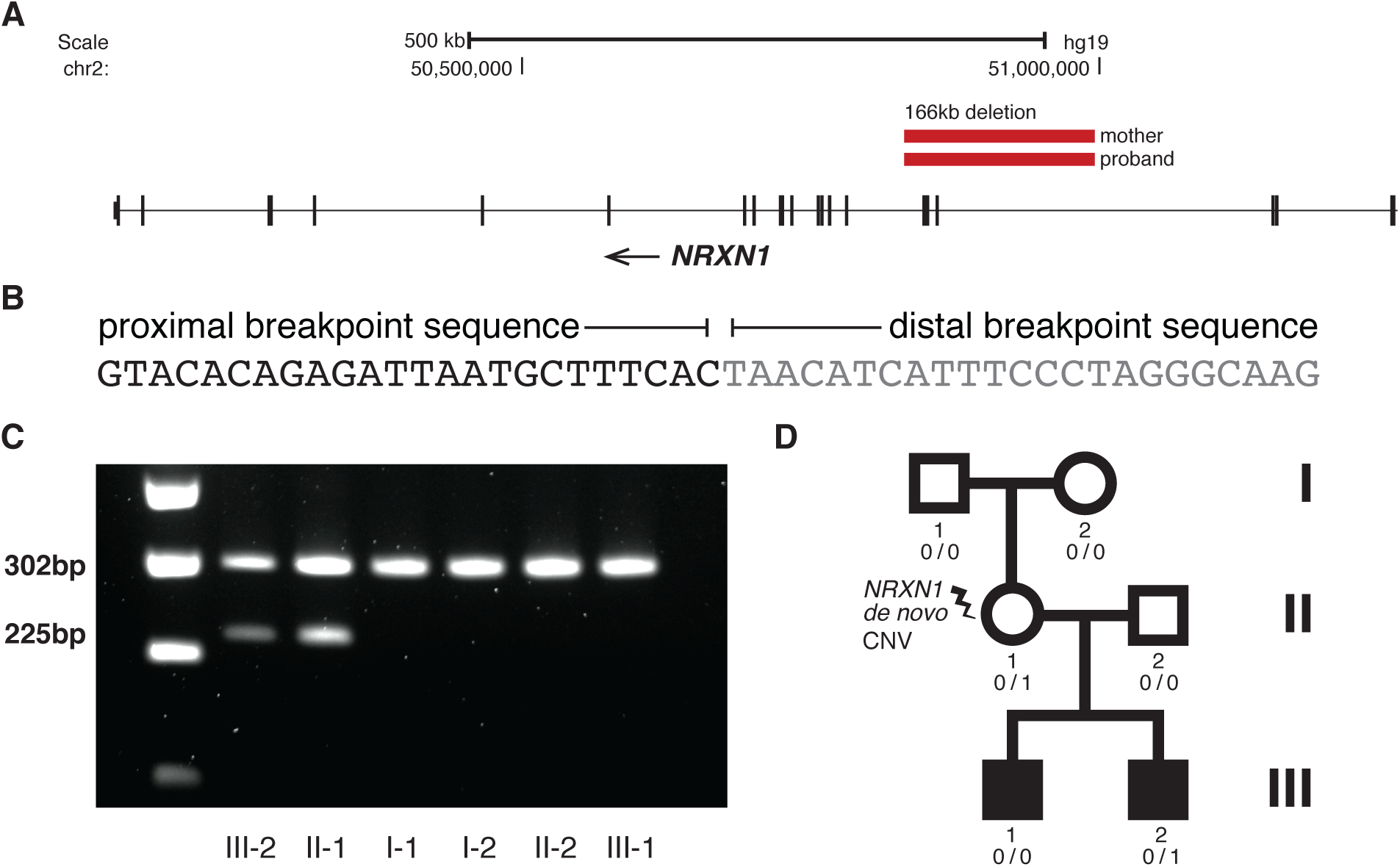
Identification and Validation of Pathogenic NRXN1 Deletion. A) 166kb deletion disrupting three exons of *NRXN1,* leading to a frameshift in the longer isoform (*α-NRXN1*). B) Breakpoint mapping shows unique sequence flanking the breakpoint, suggestive of a non-homologous end joining (NHEJ) mechanism. C) A forward PCR primer was designed proximal to the breakpoint and two reverse primers were designed, one within the deletion region that produces a 302bp product and one distal to the breakpoint, which produces a 225bp product only in the presence of a deletion. We confirm the presence of the deletion in this pedigree in the proband and mother (III-2 and II-1), but not in the father, sibling, or maternal grandparents. D) Pedigree of the multiplex-ASD family. The *NRXN1* deletion is *de novo* in the mother and passed on to her younger son. The mother is unaffected, and the older son also has ASD, but did not inherit the deletion and has less pronounced phenotypes, suggesting other *de novo* and / or inherited variants are contributing to ASD in this family.

